# A large attachment organelle mediates interaction between a novel Nanobdellota archaeon YN1 and its host

**DOI:** 10.1101/2024.05.04.592509

**Authors:** Matthew D Johnson, Hiroyuki D. Sakai, Bindusmita Paul, Takuro Nunorura, Somavally Dalvi, Manasi Mudaliyar, Doulin C Shepherd, Michiru Shimizu, Shubha Udupa, Moriya Ohkuma, Norio Kurosawa, Debnath Ghosal

## Abstract

DPANN archaea are an enigmatic superphylum that are difficult to isolate and culture in the laboratory due to their specific culture conditions and apparent ectosymbiotic lifestyle. Here we successfully isolated and cultivated a co-culture system of a novel *Nanobdellota* archaeon YN1 and its host *Sulfurisphaera ohwakuensis* YN1HA. We characterised the co-culture system by complementary methods, including metagenomics and metabolic pathway analysis, fluorescence microscopy, and high-resolution electron cryo-tomography (CryoET). We show that YN1 is deficient in essential metabolic processes and requires host resources to proliferate. CryoET imaging revealed an enormous attachment organelle present in the YN1 envelope that forms a direct interaction with the host cytoplasm, bridging the two cells. Together our results unravelled the molecular and structural basis of ectosymbiotic relationship between YN1 and YNHA. This research broadens our understanding of DPANN biology and the versatile nature of their ectosymbiotic relationships.

## Introduction

The DPANN superphylum (an acronym of the names of first 5 phylum-level groups: “Ca. Diapherotrites”, “Ca. Parvarchaeota”, “Ca. Aenigmarchaeota”, “Ca. Nanohaloarchaeota” and “Ca. Nanoarchaeota”)^1^, is a one of the most enigmatic groups in microbial ecosystems. Their metagenome-assembled genomes (MAGs) from diverse environments have been reported over the last few years, revealing that DPANNs account for approximately half of all the archaeal diversity^4,5^.

DPANN organisms are distinguished by their diminutive size, reduced genomes, and limited metabolic abilities. Therefore, the majority of the DPANNs thrive in mutualistic, commensal, or parasitic associations with a variety of archaeal and bacterial hosts^4,6–11^. Despite their widespread occurrence, and significant role in microbial ecology and environment, very little is known about their cell biology, metabolic potential, and molecular/structural basis of their symbiosis. A significant hurdle in the study of DPANNs lies in the formidable difficulty of isolating and establishing new DPANN co-culture systems. Consequently, very few DPANN co-cultures systems have been cultured and investigated to date^6,7,12–15^.

We have successfully established a new co-culture system composed of a novel *Nanobdellota* archaeon YN1 and its host *Sulfurisphaera ohwakuensis* YN1HA. We used microbiology, metagenomics and metabolic pathway analysis, fluorescence microscopy, and high-resolution electron cryo-tomography to characterise the YN1-YN1HA interaction. Our analyses revealed the growth characteristics and genomic features of YN1, and uncovered a novel attachment organelle in the YN1 envelope that bridges the host and DPANN cytoplasm, facilitating ectosymbiotic relationship.

## Results

### A novel *Nanobdellota* DPANN-host co-culture system

From an initial enrichment culture, three archaeal taxa were detected: *Sulfurisphaera ohwakuensis*^16^, *Saccharolobus solfataricus*^17^, and a novel archaeon belonging to the *Nanobdellota*^18^ (Supplementary Fig. 1A). Using the enrichment culture as a starting material, we attempted to establish a pure co-culture by dilution to extinction method^19^. After three purification attempts, a pure co-culture composed of a *Nanobdellota* archaeon (strain YN1) and *S. ohwakuensis* (strain YN1HA) was successfully established. The purity was checked by shotgun genome sequencing of genomic DNA extracted from the co-culture. As a result of hybrid assembly of Illumina short and Nanopore long reads, two complete circular genomes having 773,326 bp (for YN1) and 2,761,125 bp (for YN1HA), respectively, were obtained. No other contigs encoding any prokaryotic marker gene were obtained from the hybrid assembly.

The genome of strain YN1 has 889 protein coding sequences (CDSs), 3 rRNAs, and 43 tRNAs, with a GC content of 23.4%, whereas that of *S. ohwakuensis* YN1HA has 3,130 CDSs, 2 rRNAs, and 47 tRNAs, with a GC content of 32.6% (Supplementary table 1 and 2). Phylogenomic analysis using amino acid sequences of 53 archaeal marker genes derived from the database^20^, placed YN1 within the phylum *Nanobdellota* (Fig. 1A). The closest relative of YN1 was *Nanobdella aerobiophila* MJ1^T^, whose host was reported to be *Metallosphaera sedula* MJ1HA. This was in good agreement with the fact that YN1 and *N. aerobiophila,* but not other DPANNs, can grow under aerobic conditions. The topology of phylogenetic tree based on the 16S rRNA sequence was almost the same as that of the based on 53 archaeal marker genes (Fig. 1A and 1B). The 16S rRNA gene sequence similarity between strain YN1 and *N. aerobiophila* MJ1^T^ was only 86.5%, significantly below the highly reliable threshold value of 94.5%^23^ for defining prokaryotic genera. The growth temperature of YN1 (55-95°C) is higher than that of *N. aerobiophila* (60-75°C). In addition, the host species of YN1 is different from that of *N. aerobiophila* (i.e. *M. sedula)*. These differences indicate that strain YN1 represents a novel genus within the phylum *Nanobdellota*. Therefore, we propose a novel genus and species “*Candidatus Nanofervidus parviconus*” gen. nov. sp. nov. to accommodate strain YN1 (Na.no.fer’vi.dus. Gr. masc. n. nânos, a dwarf; L. masc. adj. fervidus, hot; N.L. masc. n. Nanofervidus, a thermophilic small-size microorganism. parviconus par’vi.co’nus. L. masc. adj. parvus, little; L. masc. n. conus, cone; N.L. fem. n. parviconus).

**Figure 1.**
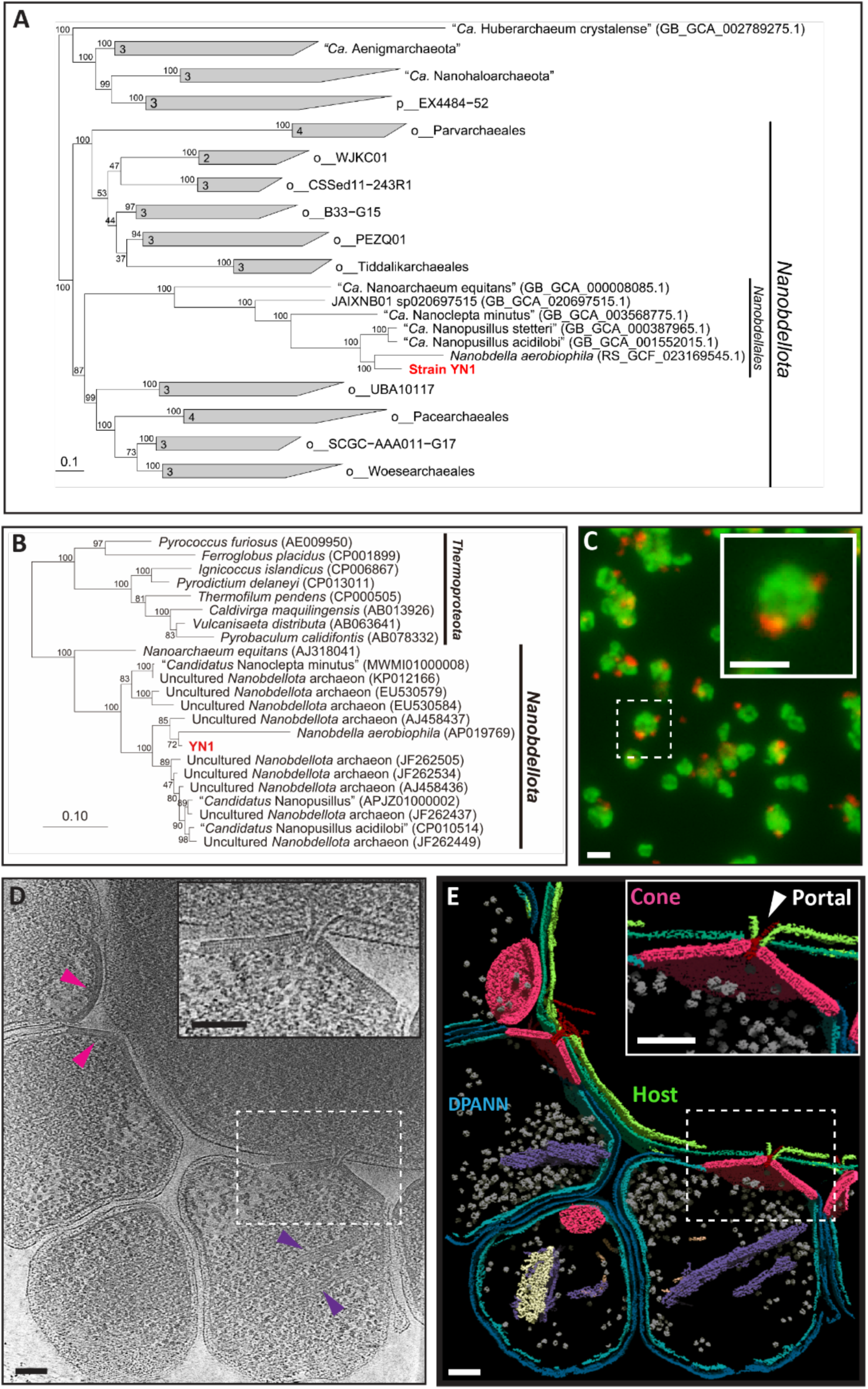
Isolation and characterization of a novel DPANN-host co-culture system. (A) The maximum-likelihood tree was reconstructed using iqtree2 with the LG+F+R6 model. The tree was reconstructed based on 53 archaeal maker protein sequences obtained from GTDB-tk (release214). The scale bar represents 0.1 amino acid substitutions per sequence position. Bootstrap values are indicated at nodes. The numbers next to the nodes at the collapsed clades are the numbers of sequences used. (B) The maximum-likelihood tree was reconstructed using iqtree2 with the GTR+F+R3 model. The tree was reconstructed based on 16S rRNA gene sequences obtained from Genbank. The scale bar represents 0.1 nucleotide substitutions per sequence position. Bootstrap values are indicated at nodes. (C) Fluorescent in situ hybridization of YN1-YN1HA co-culture. Red: *Nanobdellota* archaeon YN1, Green: *Sulfurisphaera ohwakuensis* YN1HA. (D) 2D slice through a 3D tomogram of a representative YN1-YN1HA interaction showing YN1 and YN1HA cells. Pink arrows indicate example cone structures, purple arrows indicate cytoplasmic filaments, zoomed in YN1-YN1HA interaction is shown in the inset. (E) Segmentation analysis of the volume shown in (D), YN1 cytoplasmic membrane = light blue, YN1 outer layer = dark blue, YN1HA cytoplasmic membrane = light green, YN1HA outer layer = dark green, cones = pink, cytoplasmic filaments 1 and 2 are shown in purple and white respectively, intercellular filaments = red, and ribosomes = grey. Scale bars in (D) and (E) represent 100 nm.

Growth conditions of the co-culture system were investigated to establish an optimum. The growth temperature and pH ranges for YN1 were 55-95°C and pH 2.0-3.0, respectively, while those for *S. ohwakuensis* YN1HA were 55-95°C and pH 1.5-5.5, respectively (Supplementary fig. 2A and B). The optimal conditions for both strains were established as 80°C and pH 3.0. Using these conditions, the growth of YN1 started from day 1 when its host YN1HA entered the early exponential phase (Supplementary Fig. 1B). During the middle exponential phase of YN1HA from day 1 to day 2, YN1 exhibited a growth rate almost identical to that of YN1HA. From day 2 to day 3, while YN1HA was in late exponential to early stationary phases, YN1 entered exponential phase. At this point, the growth of YN1HA significantly inhibited compared to pure culture, and many host cells were surrounded by nanosized (approximately 300 nm) cells of YN1 as observed under the scanning electron microscopy (SEM) (Supplementary fig. 1C). After day 5, when YN1 entered the death phase, the growth of YN1HA was slightly recovered and reached a maximum cell density at day 7 (Supplementary fig. 1B).

Fluorescence *in situ* hybridization (FISH) analysis confirmed that nanosized cells with approximately 300 nm in diameter were “*Ca.* Nanofervidus parviconus” YN1, while relatively larger cells with 1-2 μm in diameter were *S. ohwakuensis* YN1HA (Fig. 1C). While most YN1HA cells were surrounded by nanosized YN1 cells in the exponential phase (day 3-4), many free YN1 cells away from the YN1HA cells were also observed in fluorescently stained cells (Supplementary fig. 3A and B). Cell division of YN1 away from the host was occasionally observed under scanning electron microscopy (Supplementary fig. 1D). When YN1 cells were physically separated from the YN1HA by filtration (pore size: 0.45 μm), YN1 growth was completely inhibited. A co-cultivation screen showed that YN1 only grows on *S. ohwakuesnsis* strains as a host but not on other genera and species (i.e., *Metallosphaera*, *Sulfuracidifex*, *Sulfodiicoccus*, *Saccharolobus*, and *Sulfurishaera javensis*). Growth of co-culture was confirmed by qPCR targeting the 16S rRNA gene of each species after 1-2 weeks of cultivation. Conversely, YN1 was able to grow on different strains of *S. ohwakuensis* (YN1HA and TA-1^T^).

### CryoET uncovers an enormous attachment organelle used by YN1 to interact with its host

Electron cryo-tomography (CryoET) is unparalleled in capturing a broad range of biological phenomena from microns to sub-nanometer scale in their near-native conditions^21–24^. CryoET does not require molecular biology, and therefore can be used as an exploratory method to identify and characterize systems that are genetically intractable^25,26^. Here, we utilized CryoET to investigate the structural basis of ectosymbiosis between YN1 and YN1HA at molecular resolution. In our tomograms, the two cell types are clearly distinguishable by their size, YN1 cells measure between approx. 300 to 600 nm in diameter compared to the host cells, which measure approx. 1 to 2 micron in diameter (Fig. 1D, supplementary movie 1). Our data revealed that multiple YN1 cells interact with their host YN1HA; we observed as many as 5 YN1 per YN1HA (2.4 average YN1 per YN1HA, n=29) (Fig. 1D, Supplementary Table 2). Intriguingly, at the interface between YN1 and YN1HA, we observed an enormous macromolecular structure imbedded in the envelope of YN1 cells (Fig. 1D, pink arrows, supplementary movie 1). Analysis of these structures in segmented volumes showed that they form open-ended cone-like structures with a wide opening at the base (cytosol facing) and narrow opening at the top (extracellular facing) (Fig. 1E, supplementary movie 2). We refer to these large structures as “cones” and the narrow opening at the interface of YN1 and YN1HA as “portals”. From our 15 tilt-series, we found a total of 31 YN1 cells and 29 cone structures. All but two YN1 cells contained a cone with a maximum of 1 cone per YN1 cell (Supplementary Table 3). Our measurements of these cone structures showed that they had an average lateral length of 165.9 nm (SD = 51.4 nm, n=14), average base diameter of 288.4 nm (SD = 82.2 nm, n=14), and average height of 97.6 nm (SD = 27.6 nm, n=14) (Supplementary Table 3). We estimated the average lateral surface area of the cones at 85094.4 nm^2^ with a standard deviation of 47618.1 nm^2^, which demonstrates their enormous size and overall distribution within our dataset. We found that the cross-sectional thickness of the cones and the size of the portal were much more consistent between different cones, the average cross-sectional thickness and portal diameter was 20.8 nm (SD = 1.8 nm, n=14) and 20.4 nm (SD = 2.6, n=14), respectively. In 15 cone structures, we also observed an intercellular filament that passed from the cytoplasm of the YN1 cell to the cytoplasm of the host cell. The filaments passed through the portal of the cone and were attached to the cone via a cytoplasmic gate-like structure (Fig. 1D inset and 1E). Analysis of the segmented volume revealed that the cytoplasmic membrane of the host cell is contorted through its outer layer (S-layer) to the cone portal, creating a cytoplasmic bridge between the two cells (Fig. 1D inset and 1E inset, supplementary movie 2). Finally, in 11 of the 31 YN1 cells, we observed filament bundles in the cytoplasm. Using automated segmentation, we trained a u-net to segment these filaments, revealing two different types of intercellular filament (Fig. 1E, supplementary movie 2, purple filaments and white filaments). It is unclear if these structures are related to the cones and involved in intercellular bridging, but their measured diameters (6.6 nm, n=11) were close to those of the intercellular filaments present in the cones. Seemingly, the cone, portal, and intercellular filament together form an “attachment organelle” that facilitates the interaction between YN1 and YN1HA, and this interaction creates a cytoplasmic bridge for resource exchange.

### CryoET reveals different stages of interaction between YN1 and YN1HA

Close inspection of our tomograms revealed different stages of interaction between YN1 and YN1HA. Based on the distance between the YN1 and YNH1A, we found three distinct relationships – i) YN1 cells completely detached from their host, ii) distally attached with their host, and iii) forming a tight junction with their host. Interestingly, the detached YN1 cells still harboured the attachment organelle (cone, portal and filament) (Fig. 2A and 2B, supplementary movie 3). This suggests that attachment organelle could exist prior to interaction with the host and/or could remain intact even after YN1 is detached from their host. The distally attached YN1 cells interacted with the host by the intercellular filament structure that protrudes out of the portal from a distance of ∼40 nm (Fig. 2C and 2D, supplementary movie 4 and 5). This filament was also coated in a proteinaceous sheath which was not observed in the fully developed interaction (proximal attachment) depicted in figure 1D and E (Fig. 2C and 2D, Supplementary movie 5). Finally, we also observed YN1 cells proximally attached with the host, bridging the two cytoplasms (Fig. 1D, E, supplementary movie 2). Interestingly, in a few examples of proximal attachment, the attachment organelle seemingly did not form any cytoplasmic bridge between the two cells (Fig. 2E and 2F). The attachment organelle appeared in direct contact with the host outer-layer and the intercellular sheathed filament was visibly inside the host and extended more than 400 nm into the host (Fig. 2F, supplementary movie 5). The biogenesis of this enormous attachment organelle must require considerable resources from YN1, suggesting that this structure is critical for host-DPANN interaction and possibly resource exchange.

**Figure 2.**
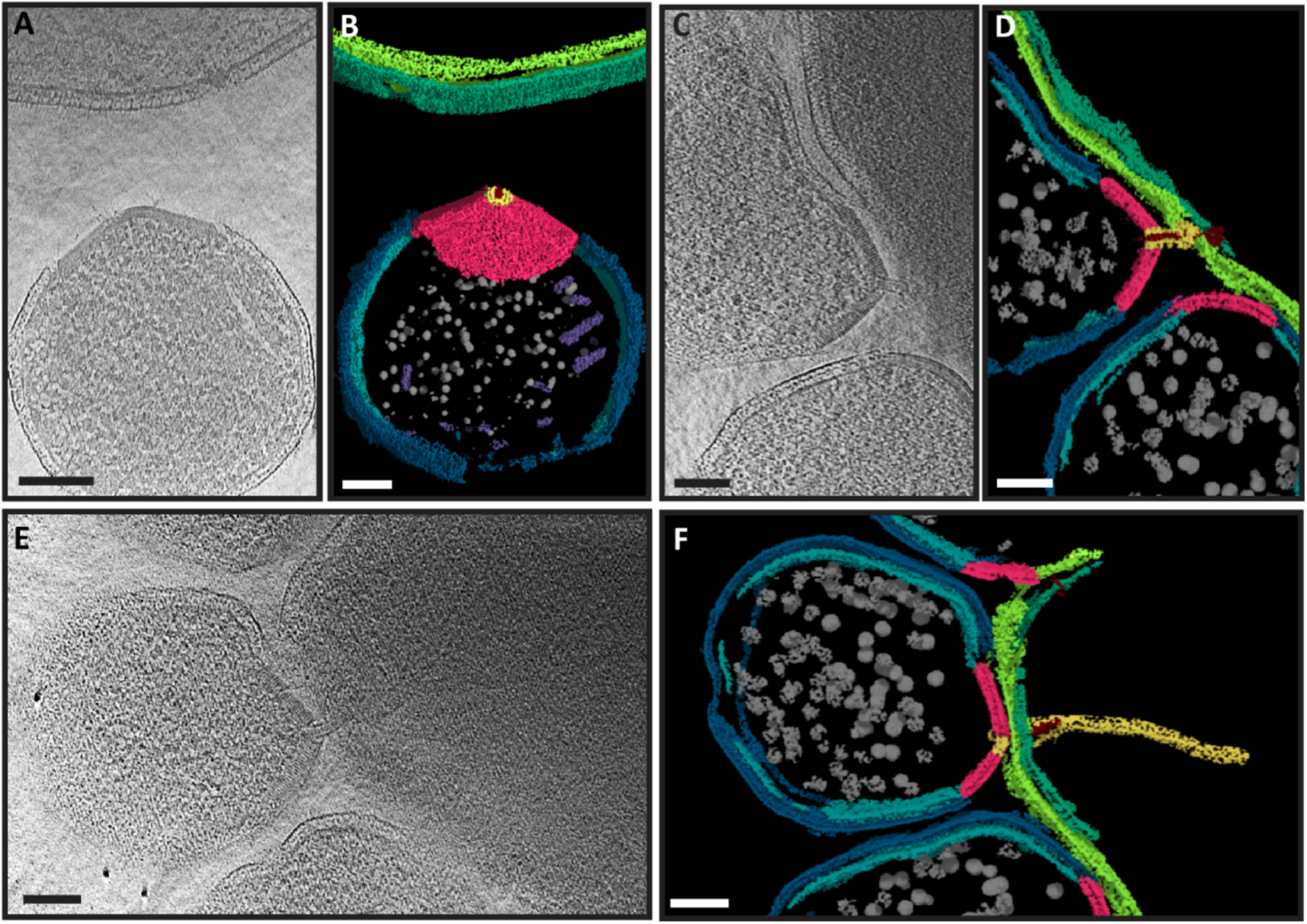
Structural basis of interaction between YN1 and YN1HA. (A) 2D slice through a 3D tomogram and (B) segmented volume of a YN1-YN1HA interaction showing separate YN1 and YN1HA cells. (C) 2D slice through a 3D tomogram and (D) segmented volume of a distant YN1-YN1HA interaction. (E) 2D slice through a 3D tomogram and (F) segmented volume of proximal/intimate YN1-YN1HA interaction. YN1 cytoplasmic membrane = light blue, YN1 outer layer = dark blue, YN1HA cytoplasmic membrane = light green, YN1HA outer layer = dark green, cones = pink, cytoplasmic filaments are shown in purple, intercellular filaments = red, intercellular sheath filament = yellow, and ribosomes = grey. Scale bars in (D) and (E) represent 100 nm.

### Metabolic pathway analysis of YN1 reveals key dependencies on the host cell

The metabolic pathways of YN1 were predicted on the basis of the genomic annotation (Fig. 3). As in the case of other *Nanobdellota* archaea, the YN1 genome lacks most of the genes in the central carbon metabolism pathways, including TCA cycle, pentose phosphate pathway, respiration, ATP synthesis, carbon fixation, and in the biosynthesis of amino acids, nucleotides, cofactors, vitamins, and lipids. Several genes related to glycolysis and gluconeogenesis were encoded in the YN1 genome; however, in glycolysis, no genes from glucose to glucose-6-phosphate (G6P), from fructose-6-phosphate (F6P) to fructose 1,6-bisphosphate (FBP), and from phosphoenolpyruvate (PEP) to pyruvate were found (Fig 3). Similarly, in gluconeogenesis, no genes from glycerate 3-phosphate (3PG) to glyceraldehyde-3-phosphate (GAP) were found. In contrast, most of the genes involved in genetic information processing (i.e., transcription, translation, replication etc.) were encoded in the YN1 genome. Kato *et al*. recently revealed the cell surface structure of *N. aerobiophila* MJ1HA^T^, and suggested it has putative S-layer proteins (MJ1_0160 and MJ1_0707)^12^. The YN1 genome also encoded these proteins (YN1_5000 and YN1_2560) with high amino acid sequence similarity to MJ1_0160 (69.5%) and MJ1_0707 (61.7%), respectively. Genes involved in archaellum formation were encoded in the YN1 genome (FlaBDEHIJF). However, under our experimental conditions, motility of YN1 was not observed under light microscopy. Genes involved in defence systems such as types I and IV of restriction and modification systems were identified. The YN1 genome did not contain the CRISPR-Cas system as there was no CRISPR repeats, and only the Cas4 gene was present.

**Figure 3.**
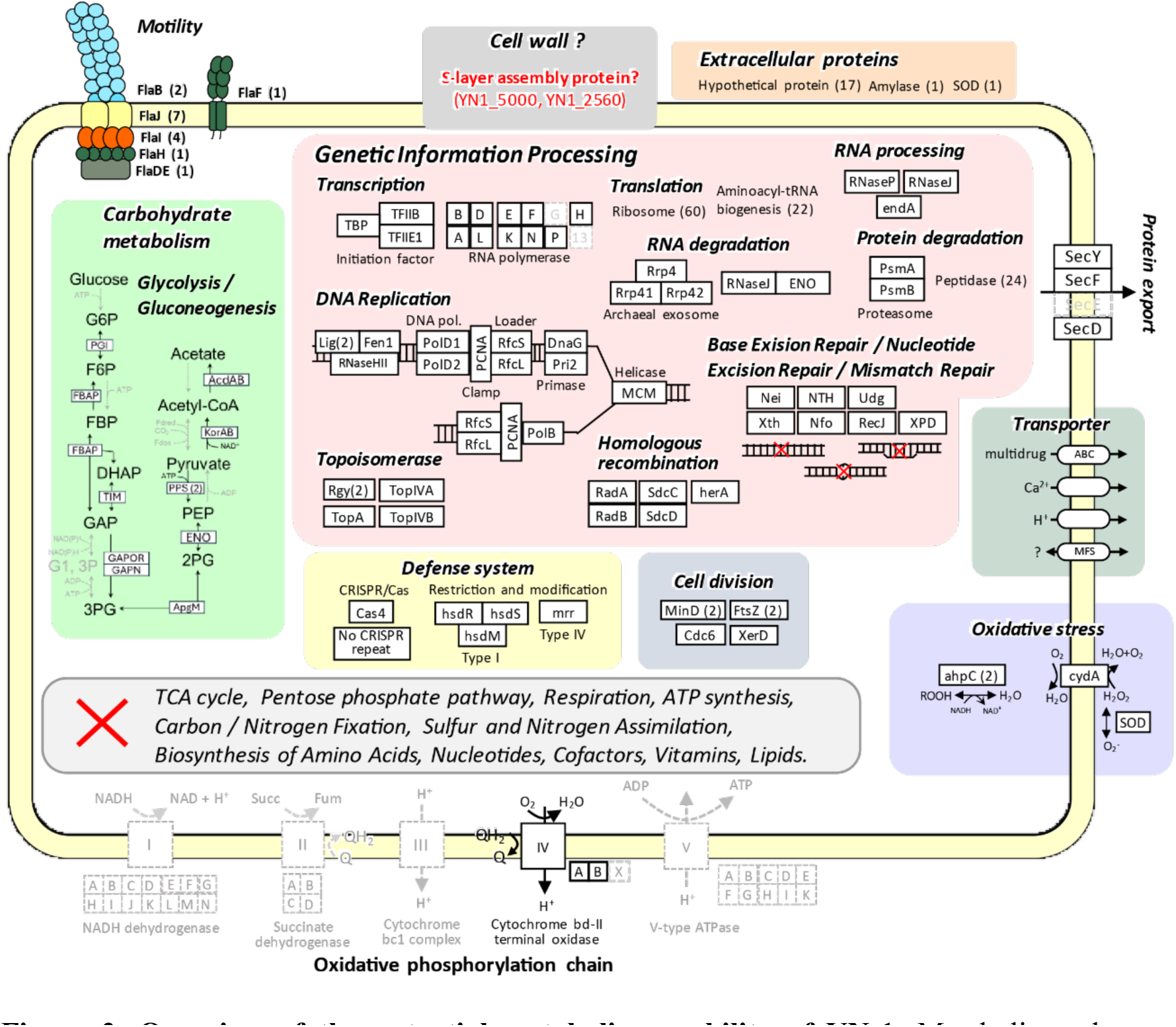
Overview of the potential metabolic capability of YN-1. Metabolic pathways were constructed based on the annotation of predicted genes (Dataset SXX). Gray indicates the lack of a gene. Coloured boxes indicate different cellular pathways. A grey box with red cross indicates pathways that are not present in YN1. The total number of genes is indicated in parentheses near the gene name.

## Discussion

The nature of DPANN biology and lifecycle is enigmatic. Based on the information of their reduced genome, DPANNs must require external resources to proliferate, but how they obtain resources and utilise them remains underexplored due to the limited number of co-culture systems established to date^6,10,12,28–30^. Previous work has shown that some DPANNs may opt for a parasitic lifestyle, instead of an ectosymbiotic relationship^28^. Furthermore, it has often been suggested that proliferation of the DPANNs would occur while attached to the host cell^4,31^. Our results also show that the YN1 genome lacks most of the genes in the central carbon metabolism, suggesting that it ought to be an obligate symbiont of other microorganisms in its natural habitat. This was also confirmed by our cocultivation experiment, where YN1 did not grow without its host, indicating that YN1 is an obligate symbiont of *S. ohwakuensis* YN1HA. The growth of *S. ohwakuensis* YN1HA significantly inhibited by the presence of YN1 (Fig. S1B), suggesting YN1HA -YN1 interaction might be parasitic. We have observed the narrow range of growth pH in YN1 (pH 2.0-3.0) compared with its host YN1HA (pH 2.0-5.5). A similar observation was reported for *N. aerobiophila*^18^, suggesting that this characteristic is a common feature in *Nanobdellota*. The presence of most genes involved in genetic information processing and the observation of dividing YN1 cells while detached from their host (Supplementary Fig. 1D) imply that YN1 has the capability to undergo cell division away from the host cell. Homologous genes for the recently suggested putative S-layer proteins of *Nanobdellota*^12^ were found in the YN1 genome (YN1_5000 and YN1_2560), suggesting that these genes may be involved in S-layer formation. The presence of genes involved in archaellum formation^32^ (FlaBDEHIJF) and the observation of archaellum-like structures in our micrographs suggest the potential for motility in YN1 (Supplementary Fig. 1E). Under our experimental conditions, however, we failed to observe motility of YN1 under light microscopy, it is possible that specific required conditions were not met for motility during our experiment.

Our co-cultivation experiments confirmed that YN1 grows on different strains of *S. ohwakuensis* (YN1HA and TA1^T^) but not on the other genera and species examined so far. (*M. sedula* DSM5348^T^, *M. hakonensis* DSM7519^T^, *M. cuprina* JCM15769^T^, *Sulfodiicoccus acidiphilus* HS-1^T^, *Sulfuracidifex tepidarius* JCM16833^T^, *Saf. metallicus* DSM6482^T^, *Saccharolobus solfataricus* JCM8930^T^, *Scl. shibatae* DSM5389^T^, *Scl. caldissimus* HS-3^T^, *Sulfolobus acidocaldarius* DSM639^T^)^17,33–38^. This is consistent with the published literature - all the cultivated *Nanobdellota* archaea reported to date can only use a specific host^9,18,39–41^. For instance, a recently described *N. aerobiophila* MJ1^T^ showed that it only grows with the original host, *M. sedula* MJ1HA, but not with the type strain of *M. sedula* TH2^T^ as well as other several different genera and species^18^. A similar observation was also reported for “*Ca. Nanoarchaeum equitans*”^41^, the first cultivated species in the phylum *Nanobdellota*^9^. In contrast, YN1 was able to grow with different strains of *S. ohwakuensis*, a unique observation in phylum Nanobdellota. This suggests that this taxon could have a narrow range of host specificity. The fact that the YN1 cells passed through the membrane filter grew with different strains of *S. ohwakuensis* possibly indicates that the free YN1 cells observed under microscopy are living cells and capable of proliferation (Supplementary Fig. 1E). The archaellum, typically known for its functions in chemotaxis and surface attachment^32^, could in principle support free YN1 cells in locating and reattaching to host cells.

We identified an enormous attachment organelle in the YN1 cell envelope that is used to attach to the YN1HA host cell to form a cytoplasmic bridge. This organelle was comprised of a cone structure with a portal opening, and a filament extension that passes through the portal. When YN1 made intimate contact with the host (proximal attachment), the intercellular filament traversed inside the host and appeared to have a sheath. Similar cone-shaped protein complexes have been previously observed in other microorganisms e.g., virus-associated pyramids (VAPs) were described in *Sulfolobus islandicus*^42^. The VAP structure did not have a portal and, although the inner layer of the VAP formed a continuous density with the inner membrane, the VAP was not embedded within the cell envelope as reported in this study. Archaeal cones have also been described in *Thermococcus kadokaraensis*^26^ and *Pyrococcus furiosus*^43^ and were described as organisation centre for archaellum assembly. However, these cone structures were cytoplasmic and without a portal. The attachment organelle of YN1 is embedded within the envelope, has filaments passing through the cone’s portal and is therefore, structurally unique. Although it is possible that these previously described structures are evolutionarily linked to the cone structures observed here, the structures are considerably different and likely have completely different functions. In one instance attachment organelles from different YN1 cells appear to be facing and possibly interacting with each other (Supplementary Fig. 4A and B, SI movie 6). However, the portals of the attachment organelles are not in plane and there is no density connecting them.

Our results show that YN1 *Nanobdellota* assembles an attachment organelle to form direct cytoplasmic bridge with the host cell. We have also described attachment organelles in YN1 cells that are detached from the host, either primed to form an interaction or a relic from a previous interaction. Finally, we described two further attachment stages where the filament, and filament sheath, of the attachment organelle appear to be inserted into the host cytoplasm, while the cone structure remained distally connected. We predict that YN1 has a biphasic lifestyle, an obligate ectosymbiont that uses its host to acquire the essential nutrients for growth, but once fuelled up, YN1 can detach and proliferate in isolation. We propose that the attachment organelle is used to latch onto the host cells and form an intimate interaction that results in the formation of a cytoplasmic bridge between the host cell and YN1, allowing for resource exchange. Many steps in this process remain unanswered, the protein(s) that comprise the attachment organelle are unknown, as is the mechanism by which they assemble this massive macromolecular complex. Interestingly, we identified different types of filaments (and filament bundles) present in the DPANN cytoplasm, but further investigation is required to confirm their identity and functions (Supplementary Fig. 4 A-E).

Our work here established a new *Nanobdellota* co-culture system and characterised the structural basis of host-DPANN interaction through complex attachment organelle. The proteins that comprise this complex remain unknown and are the subject of future research, as is the method by which this enormous organelle assembles into the YN1 membrane, permeates the host’s outer layer and fuses with the cytoplasm. This research broadens our understanding of DPANN biology and reveals the versatile nature of their ectosymbiotic relationships.

## Acknowledgements

This project was funded by an NHMRC grant (APP1196924 to DG), an HFSP grant (RGEC33/2023) to DG and HDS, The Japan Society for the Promotion of Science through Grants-in-Aid for Scientific Research (21K15153 to HDS). We thank the Ian Holmes Imaging Centre at the University of Melbourne for access to microscopes.

## Data availability

Raw read data of genome sequencing have been deposited in DDBJ/ENA/GenBank under the accession numbers DRR287339 and DRR287338. Genomic sequences of YN1 and Sulfurisphaera ohwakuensis YN1HA were deposited under the accession numbers AP031373 and AP031374, respectively.

## Materials and Methods

### Sampling

Approximately 50 mL of hot water samples containing a small quantity of mud were collected in polypropylene tubes from two sites at an acidic hot spring Yunoike in Kirishima geothermal area located in Kagoshima prefecture, Japan (Fig. S1). Temperature and pH at the sampling site were 78°C and pH 2.7, respectively (Urayama et al. 2024). The sample in a plastic tube was transported to the laboratory under ambient temperature and used for enrichment culture.

### Enrichment culture

Enrichment culture was conducted using modified Brock’s basal salt medium^16^ supplemented with 0.1% (w/v) yeast extract (MBSY medium). The pH of the medium was adjusted to 2.9. A 100 μL of each sample was inoculated into a 10 mL MBSY medium in a 20 ml screw-capped test tube (16.5 × 105 mm, ST-16.5L, Nichiden-rika glass, Kobe, Japan) and incubated on static condition at 80°C. Microbial growth of the enrichment cultures was regularly monitored by measuring the optical density at 600 nm. When the growth entered the stationary phase, a 2 mL of each enrichment culture was centrifuged (20000 x g, 25°C, 30 min) and the resulting pellet was used for DNA extraction for 16S rRNA gene amplicon sequencing. The rest of the enrichment culture was stored in a freezer at -80°C with 10% (v/v) glycerol as a cryoprotectant reagent. The cryostock was later used for the establishment of pure-co-culture composed of a *Nanobdellota* archaeon (strain YN1) and its host *Sulfurisphaera ohwakuensis* (strain YN1HA).

### 16S#rRNA gene amplicon sequencing

Microbial DNA from the enrichment culture was extracted using Extrap Soil DNA Plus Kit version 2 (Nippon Steel and Sumikin Eco-Tech). The extracted DNA was used for the 16S rRNA gene amplicon sequencing. The V3-V4 region of 16S rRNA was amplified using the DNA polymerase KOD Fx Neo (TOYOBO) with primers A340F^44^ and 806rB^45^ (with barcode for Illumina MiSeq sequencer). The following thermal cycler protocol was used: an initial denaturation of 2 min at 94°C, 25 cycles of 10 seconds denaturation at 98°C, 30 sec annealing at 50°C, followed by 68°C for 15 sec, and a final extension of 68°C for 5 min. The PCR product was sent to a sequencing company (Fasmac, Kanagawa, Japan), and the 16S rRNA gene amplicon sequencing was carried out using Illumina MiSeq sequencer (300 bp x 2). The microbial community structure of the enrichment culture was investigated by the QIIME2 pepeline^46^, followed by BLASTN search against nt database in NCBI.

### Primer design

The specific primers targeting the 16S rRNA gene for a *Nanobdellota* archaeon (NanoF: GGCGAAATGCAGTAATCCCG, NanoR2: TAACGGCTTCCCTATCCCAC) detected in the enrichment culture were designed by using Primer BLAST^47^, while those for other detected archaea (*Sulfolobales* species) were designed as previously described^30^.

### Establishment of pure co-culture and isolation of pure host

To establish a pure-co-culture only composed of a *Nanobdellota* archaeon and its host, dilution to extinction method was carried out three times under the same condition as enrichment culture. The presence of a *Nanobdellota* archaeon was confirmed by qPCR with specific primers. The same PCR cycling conditions as described previously^30^ were used. A pure-culture of host was isolated by single colony isolation from the pure-co-culture using a MBSY plate medium solidified with 0.7 % (w/v) gellan gum (Wako), 10 mM MgSO4 and 2.5 mM CaCl2.

### Genome sequencing and assembly

Microbial cells were collected by centrifugation (15,880 × g, 10 min) from the pure co-culture. Genomic DNA was extracted from the cells using Genomic-Tip 100/ G (QIAGEN). To obtain long reads, a DNA library was prepared following the protocol as described by Oxford Nanopore Technologies (NBE_9065_v109_revZ_ 14Aug2019), followed by sequencing with MinION sequencer using R9 flow cell (Oxford Nanopore Technologies). To obtain short reads, the genomic DNA was sent to Novogene Bioinformatics Technology (China) to be sequenced on an Illumina NovaSeq 6000 platform (2 x 150 bp). The short and long reads were quality filtered with using the same protocol as described previously^48^ and coassembled using Unicycler (ver. 0.4.8)^49^.

### Genome analysis

Annotation of genome sequences was carried out using Prokka (ver. 1.14.6)^50^, DFAST^51^, RAST server^52^, Kyoto Encyclopedia of Genes and Genomes (KEGG) pathway tools^53^, eggNOG-mapper (ver. 2.1.12)^54^, COGclassifier (https://github.com/moshi4/COGclassifier), and hmmscan against MEROPS^55^. The annotated data obtained by these programs were manually curated. Subcellular localization of each protein encoded in the genome sequences was predicted using PSORTb 3.0.314. The 16S rRNA gene sequence similarity between YN1 and its most related species *Nanobdella aerobiophila* MJ1^T^ was calculated by BLASTN whereas their average amino acid identity (AAI) was calculated using the Kostas laboratory server (enve-omics.ce.gatech.edu/ani/).

### Phylogenomic analysis

To conduct the phylogenomic analysis of strain YN1, amino acid sequences of 53 archaeal marker genes defined by the GTDB database (release 214)^20^ were collected using GTDB-Tk (ver. 2.3.0)^56^. The collected sequences were realigned by MUSCLE^57^, followed by trimming using TrimAL^58^ with automated1 option. A maximum likelihood phylogenetic tree was reconstructed using IQ-TREE2^59^ with the LG+F+R6 model. Bootstrap support values were calculated with 1,000 replicates.

### Growth monitoring

The growth of a *Nanobdellota* archaeon YN1 and its host *Sulfurisphaera ohwakuensis* YN1HA in pure-co-culture or pure culture was monitored by qPCR with specific primers for each species. The same protocols as described previously^30^ was used. In brief, microbial DNAs regularly extracted from cultures were applied to qPCR with specific primers targeting 16S rRNA gene, which is a single copy gene for each strain. Cultivation was conducted in triplicate to compare YN1HA growth between co-culture and pure culture, and in duplicate to examine the growth temperature and pH ranges for YN1 and YN1HA.

### Light microscopy and Scanning electron microscopy

General cell morphology was assessed by differential interference contrast microscopy (BX51; Olympus). The cells stained with SYBR Green I by the same protocol as described previously^30^ were observed by fluorescent microscopy (BX51; Olympus). Scanning electron microscopy (JSM-7500F; JEOL) was carried out using the same protocols as described previously^30^.

### Fluorescent in situ hybridization

Fluorescent in situ hybridization (FISH) was conducted using the method as previously described^60^ with some modifications. The FISH probes used for YN1 (Nano-R4_TEX: GTATTCCCGTGGCGACTGC) and YN1HA (SFB-R_FAM: CGGTTACTAGGGATTCCTCG), both targeting the 16S rRNA gene were designed using Primer BLAST^47^. The probes were labelled with either Texas-red or 6-carboxyfluorescein at their 5’ end. Cells were fixed with 2% paraformaldehyde for overnight, washed in phosphate buffered saline (PBS), and resuspended in PBS before spotting 15 µL onto a silane-coated glass slide (S8111; MATSUNAMI). The spotted sample was dried, incubated in 100 µl of 0.25 N HCl solution for 30 minutes at room temperature, rinsed in distilled water, and dehydrated by serial treatment with 50%, 80%, 90%, and 100% ethanol solutions (3 minutes each, 2 times). Hybridization was carried out with the probes (final concentration: 1 pmol/µl each) in buffer (0.9 M NaCl, 20 mM Tris-HCl, 0.1% SDS, pH 7.5) at 48°C overnight. After hybridization, the slide was washed in buffer (0.9 M NaCl, 20 mM Tris-HCl, 0.1% SDS, 5 mM EDTA, pH 7.5) at 48°C for 20 min, lightly rinsed in distilled water, and enclosed with mounting medium (SlowFade Gold Antifade reagent; Life Technologies). The fluorescence signals were detected with an epifluorescence microscope (BX63; Olympus) fitted with filter sets specific for Texas-red and fluorescein.

### Cocultivation experiments of YN-1 with various thermoacidophilic species

To investigate the host specificity of YN1, the following thermoacidophilic strains were cocultivated with YN1*: M. sedula* DSM5348^T^, *M. hakonensis* DSM7519^T^, *M. cuprina* JCM15769^T^, *Sulfodiicoccus acidiphilus* HS-1^T^, *Sulfuracidifex tepidarius* JCM16833^T^, *Saf. metallicus* DSM6482^T^, *Saccharolobus solfataricus* JCM8930^T^, *Scl. shibatae* DSM5389^T^, *Scl. caldissimus* HS-3^T^, *Sulfolobus acidocaldarius* DSM639^T^, *Sulfurisphaera. ohwakuensis* TA-1^T^, *S. ohwakuensis* YN1HA (original host), and *S. javensis* KD-1^T^. These strains were obtained from the German Collection of Microorganisms and Cell Cultures (DSMZ) or Japan Collection of Microorganisms (JCM), except for *Sfd. acidiphilus* HS-1^T^, *Scl. caldissimus* HS-3^T^, *S. ohwakuensis* TA-1^T^, and *S. javensis* KD-1^T^, that were originally isolated and stored in our laboratory (Soka University). They were cultivated in MBSY medium (pH 3.0), with the exception of *Saf. tepidarius* JCM16833^T^ and *Saf. metallicus* DSM6482^T^, that were cultivated in MBSY medium with 1 g/L elemental sulfur. Pure YN1 cells were collected from the original pure co-culture (YN1+YN1HA) by filtration using a sterilized syringe filter (pore size: 0.45 μm; material: polyethersulfone). A 60 μL of each pure culture and 600 μL of the filtrate containing YN1 cells were inoculated into 6 mL of cultivation medium in a 20 ml screw-capped test tube (16.5 × 105 mm, ST-16.5L, Nichiden-rika glass, Kobe, Japan). As a negative control, 1 mL of the filtrate was inoculated into 5 mL of MBSY medium without the inoculation of any of the strains. All these cultivations were conducted in duplicate. The incubation temperatures were set at 65°C for genera *Metallosphaera*, *Sulfodiicoccus*, and *Sulfuracidifex*, 75°C for *Scl. solfastricus*, *Scl. shibatae*, *Slf. acidocacldarius*, and 80°C for *S. ohwakuensis* YN1HA, *S. ohwakuensis* TA1^T^, and *Scl. caldissimus* HS-3^T^ (under aerobic conditions). At the stationary growth phase, the growth of YN1 was assessed by qPCR with specific primers.

### Sample vitrification and Cryo-ET data acquisition

Before vitrification, archaeal co-cultures were mixed in a 2:1 ratio with 10 nm colloidal gold beads precoated with 1% BSA (Sigma-Aldrich, Australia). The mixture was added onto glow-discharged copper R2/2 Quantifoil holey carbon grids (Quantifoil Micro Tools GmbH, Jena, Germany). Grids were blotted for 6-8 seconds (under 100% humidity conditions) and plunged into liquid ethane using a Vitrobot Mark IV, FEI (Thermo Fisher Scientific). Micrographs of YN1 co-cultures were acquired using an FEI Titan Krios G4, 300 keV FEG transmission electron microscope (Thermo Fisher Scientific), equipped with a BioQuantum K3 Imaging Filter (slit width 20 eV), and a K3 direct electron detector (Gatan). Tilt series were collected automatically using FEI Tomography 5 from -60° to +60° at 2° intervals with a defocus of -6 μm and a total electron fluence of 120 e^-^/Å^2^ and a pixel size of 3.4 Å.

### Tomogram reconstruction

Before processing of tilt series, tilt images of poor quality were identified and removed through visual inspection. The cleaned tilt series were aligned using fiducial tracking in IMOD (Version 4.11.5) ^61,62^ and subsequently binned 4 times (13.56 Å/pix). Tomo3D (version 2.2)^63^ was used to generate SIRT reconstructed tomograms from aligned tilt series, which were then filtered to enhance contrast using the deconvolution filter in IsoNet^64^.

### Tomogram Segmentation and Visualization/ 3D-segmentation

Visualization and segmentation of tomograms were conducted using the Dragonfly software (https://www.theobjects.com/dragonfly/index.html). Tomograms were preprocessed using built-in filters, including histogram equalization, Gaussian, and unsharp filters. Neural network 5-class U-Net (with 2.5D input of 5 slices), were trained on tomogram slices to recognize YN1 cytoplasmic membrane, YN1 outer layer, YN1HA cytoplasmic membrane, YN1HA outer layer, cones = pink, cytoplasmic filaments, intercellular filaments, intercellular sheath filament and ribosomes and further fine-tuned to achieve high-quality segmentation for automated 3D segmentation. The software’s built-in tools for exporting 2D images and creating 3D movies were utilized.

---

**Supplementary figure 1.**
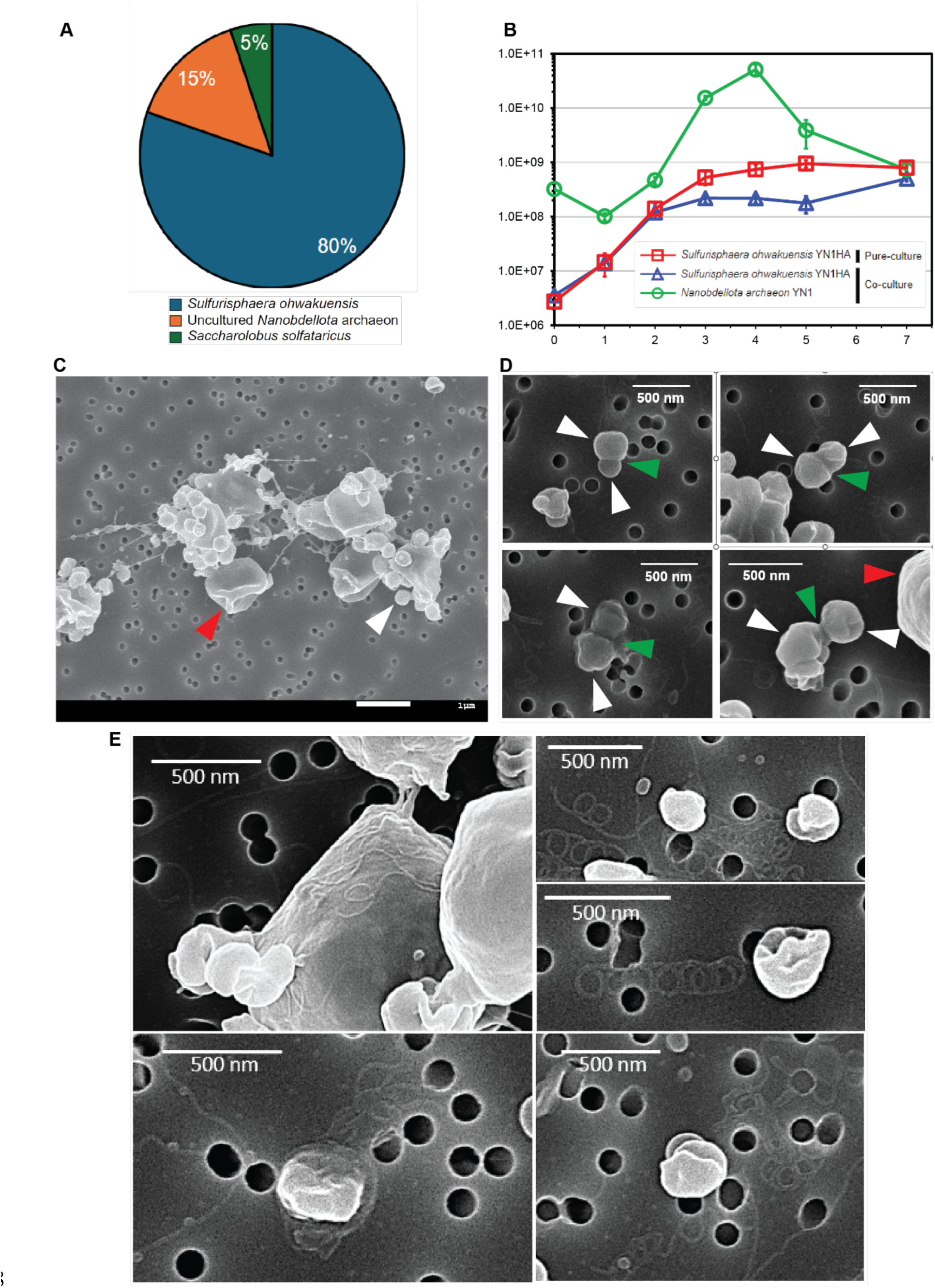
The YN1 and YN1HA co-culture system, metagenomics, growth and association. **(A)** Pie chart depicting the archaeal community structure of initial enrichment culture, based on the 16S rRNA amplicon analysis. **(B)** Growth curve analysis of pure YN1HA and in co-culture with YN1 using 16S-rRNA quantification. **(C)** SEM analysis of an example co-culture used in this work showing small YN1 cells attached to larger YN1HA cells. **(D)** SEM analysis showing example cell division events of YN1 cells separate from the host cell. **(E)** SEM analysis showing the presence of archaellum both on the host cell (top left panel) and in the extracellular milieu (right side and bottom left panel). Red, white, and green arrows indicate YN1HA cells, YN1 cells, and YN1 cell division sites respectively.

**Supplementary figure 2.**
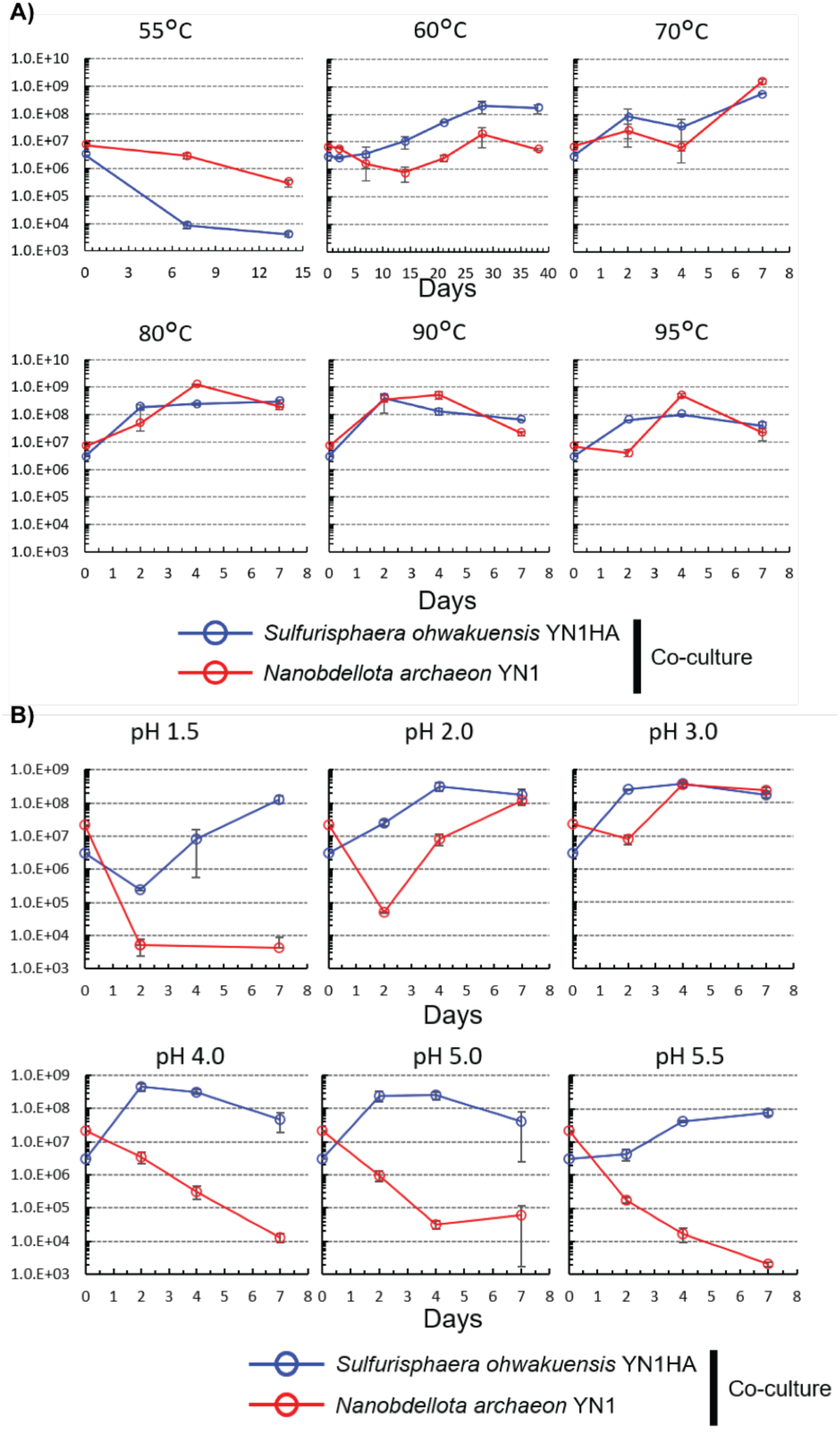
Optimal growth conditions for the YN1 – YN1HA co-culture system. **(A)** growth curves of YN1 and YN1HA at different temperatures. **(B)** Growth curves of YN1 and YN1HA at different pH.

**Supplementary figure 3.**
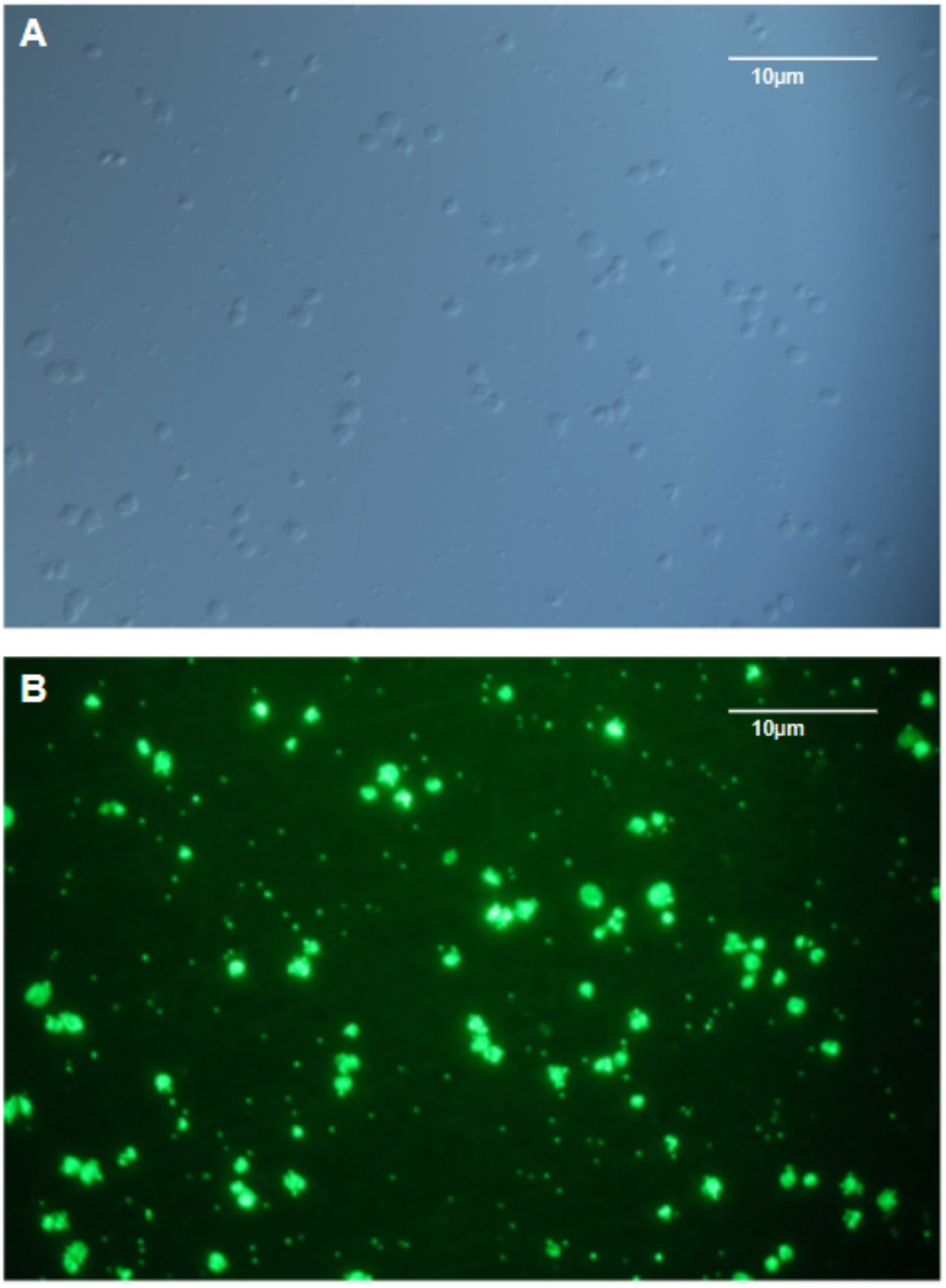
Differential interference contrast microscopy. **(A)** and fluorescence microscopy (BX51, Olympus) **(B)** of YN1-YN1HA co-culture stained with SYBR green.

**Supplementary figure 4.**
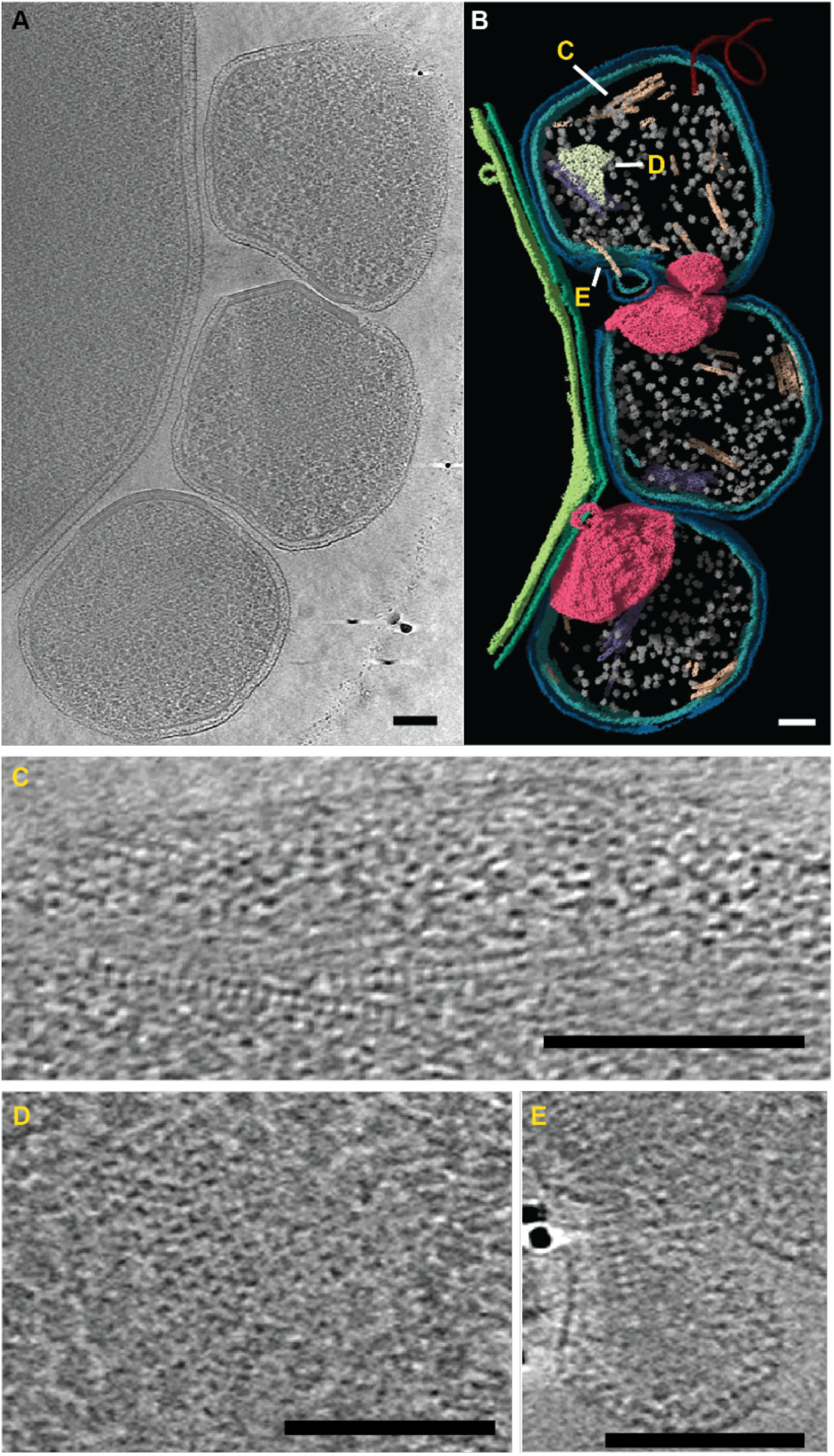
Additional features in YN1 cells. **(A)** 2-D slice through a 3-D tomogram showing a host interacting with three YN1 cells. **(B)** Segmented volume of the same tomogram as in (A), highlighting interesting features. (C-E) 2-D slices from the same tomogram shown in (A) focussing on the interesting features, **(C)** striated filaments, **(D)** 2D matrix, **(E)** filaments inside a membrane protrusion. Scale bars represent 100 nm.

## Supplementary Information Movie legends

**SI Movie 1**

Tomographic movie moving through the z axis showing the interaction of YN1 DPANN cells to their YN1HA host cell. Also shown are the intercellular and cytoplasmic filaments.

**SI Movie 2**

Three-dimensional (segmented) view of an YN1 and YN1HA interaction showing examples of the attachment organelles connecting the two cell types. Also shown are the intercellular and cytoplasmic filaments.

**SI Movie 3**

Tomographic movie moving through the z axis showing the interaction of YN1 cells forming various interactions with a YN1HA host cell. Also shown are the intercellular, cytoplasmic, and sheath filaments

**SI Movie 4**

Three-dimensional (segmented) view of a YN1 cells forming various interactions with a YN1HA host cell. Also shown are the intercellular, cytoplasmic, and sheath filaments.

**SI Movie 5**

Tomographic movie through the z axis of a YN1 cell searching for a host, showing the presence the attachment organelle before host cell interaction.

**SI Movie 6**

Tomographic movie moving through the z axis showing two YN1 cells in proximity with their attachment organelles forming the interacting surfaces.

## Supplementary Tables

**Supplementary table 1**

YN1 genome annotation.

**Supplementary table 2**

YN1HA genome annotation.

**Supplementary table 3**

Quantification of features observed in CryoET tomograms.

